# A human cytotrophoblast-villous endothelium-fetal organ multi-cell model and the impact on gene and protein expression in placenta cytotrophoblast, fetal hepatocytes and fetal kidney epithelial cells

**DOI:** 10.1101/2025.04.01.646643

**Authors:** Helen N. Jones, Rebecca L. Wilson

**Author notes:** **Corresponding Author:** Dr. Rebecca L. Wilson, Center for Research in Perinatal Outcomes, University of Florida, Gainesville, Florida 32610.

## Abstract

Appropriate fetal growth during pregnancy requires multi-directional communication from the maternal, placental and fetal systems. Disruption in any of these signaling arms can have deleterious consequences for fetal growth and initiate developmental adaptations within fetal tissues and organs that are associated with both short- and long-term morbidities. In this proof-of-concept translational, human cell model study we aimed to identify the impacts of altered trophoblast stress response mechanisms and human *insulin-like 1 growth factor* (*hIGF1*) nanoparticle gene therapy on gene and protein expression in fetal liver hepatocytes and fetal kidney epithelial cells. We utilized human cell lines: BeWo choriocarcinoma cells (trophoblast), Human Placental Micro-Vascular Endothelial Cells, and WRL68 (hepatocytes) or HEK293T/17 (kidney epithelium), in a co-culture model designed to mimic cytotrophoblast-villous endothelium-fetal organ communication. Trophoblast stress response mechanisms were increased by culturing BeWo cells in growth media without FBS. Stressed BeWo cells were also treated with a *hIGF1* nanoparticle gene therapy known to mitigate cellular stress mechanisms. Stressed BeWo cells had increased expression of cellular stress mechanisms but not when *IGF1* was over-expressed with a transient *hIGF1* nanoparticle gene therapy. Stressed and Stressed+*hIGF1* BeWo cells had increased expression of gluconeogenesis and glycolysis rate-limiting enzymes. Gene and protein expression in fetal liver and kidney cells was not impacted by increased trophoblast stress or *hIGF1* nanoparticle gene therapy. In conclusion, our data demonstrated that cytotrophoblast under stress turn on mechanisms involved in glucose production. Whether this is reflected in vivo remains uninvestigated but may represent a placental compensation mechanism in complicated pregnancies.

## Introduction

Advances in epidemiological research and animal studies have illuminated a significant association between the in utero environment and the likelihood of developing diseases, such as diabetes and cardiovascular disease, later in life [1, 2]. This association fundamentally alters the idea that a person’s vulnerability to diseases solely arises from genetic and postnatal environmental interactions. The Developmental Origins of Health and Disease (DOHaD) hypothesis predominantly links fetal growth restriction (FGR), as a result of placental insufficiency and inadequate nutrient and oxygen delivery to the fetus, with increased risk of developing physiological deficits including glucose intolerance, insulin resistance and hypertension, as early as adolescence [3]. However, there is emerging evidence that fetal overgrowth, as a consequence of excessive nutrient supply to the fetus, can also have a profound impact on lifelong health and disease [4]. These scenarios ultimately underscore the critical role of placental function and fetal nutrient status in fetal developmental programming and shaping long-term health outcomes.

Appropriate in utero fetal growth requires the placenta to delicately balance fetal demands with maternal supply [5]. The placenta functions as an endocrine organ that coordinates the transfer of nutrients and waste between mother and fetus and facilitates gas exchange [6]. Nutrient transfer specifically, occurs via facilitated diffusion (glucose and lactate), active transport with the aid of carrier proteins, endocytosis and/or exocytosis and increases as the fetal growth rate increases [7]. The functional area for nutrient transfer in the placenta is the chorionic villous tissue, which contains the placental blood vessels that connect directly to fetal circulation via the umbilical cord [8]. Chorionic villi are covered in a multinucleated syncytium which is in direct contact with the maternal blood within the intervillous space. Underneath this syncytial layer lie the cytotrophoblast cells, placental macrophages, fibroblasts and capillary endothelium containing the fetal blood [9]. Hence, nutrients must cross several layers of cells and basement membranes in order to move between maternal and fetal circulations.

It has long been understood that maternal-placental-fetal communication is multi-directional, particularly when it comes to fetal growth [5]. The fetus communicates with the mother via the placenta which in turn produces the hormones and factors necessary to promote appropriate maternal metabolism and behaviors but may be constrained depending on the maternal state. Hence, dysfunction in any of these signaling arms can lead to alterations in placental function and fetal growth, ultimately initiating a cascade of developmental adaptations with adverse consequences both in the short and long term. Placental secretion of proteins including hormones [10], cytokines and growth factors [11], as well as glycoproteins, steroid hormones and more recently extracellular vesicles [12], have been heavily studied to better understand their implications to maternal pregnancy adaptation as well as fetal developmental programming of adverse long-term health. For example, insulin-like 1 growth factor (IGF1) is a growth factor produced by the placenta from the first trimester of pregnancy [13]. IGF1 is needed for appropriate placental growth and development and influences transport of nutrients including glucose and amino acids across the placenta [14]. In humans, both maternal and fetal circulating levels of IGF1 are lower in cases of FGR compared to appropriate for gestational age cases [15, 16]. Transgenic mouse models have also confirmed the necessity for placental *Igf1* in the promotion of fetal growth [17, 18].

Whilst there are many extrinsic and intrinsic causes of perturbed fetal growth, the commonality between the causes is placental dysfunction [19-21]. Hence, therapeutic interventions which target the placenta offer unparalleled opportunities to not only prevent adverse maternal and fetal outcomes in the short-term perinatal period but also mitigate the risk of major diseases throughout an individual’s lifespan. Whilst prior investigations into the use of IGF1 as a potential therapeutic intervention have had mixed results [22-24], we have been successful in showing that specifically increasing placental gene expression of human *IGF1* (*hIGF1*) maintains or mitigates against reduced fetal weight in various animal models [25-27]. Placental gene expression of *hIGF1* is increased with a plasmid that contains the *hIGF1* gene under the control of a placenta-specific promotor (*PLAC1* or *CYP19a1*), with cellular uptake aided by the use of a non-viral polymer nanoparticle [26, 28, 29]. Our animal studies are supported by in vitro investigations in human placenta models which confirm the ability to manipulate cytotrophoblast nutrient transporter expression and prevent increased cell death under oxidative stress condition by treatment with the *hIGF1* nanoparticle [29, 30].

In addition to comprehensively characterizing the placental response to *hIGF1* gene therapy, we also understand that manipulating the placenta indirectly impacts fetal liver and kidney development and function [31]. More specifically, improving development and function with *hIGF1* gene therapy mitigates FGR-associated changes in gene and protein expression of factors relating to glucose metabolism in the fetal liver [31] and blood pressure regulation in the fetal kidneys [32]. Overall, suggesting the potential to prevent or reverse developmental programming of metabolic health deficits with an in utero intervention. In this proof-of-concept translational, human cell model study we aimed to identify the impacts of altered trophoblast stress response mechanisms and *hIGF1* nanoparticle treatment on gene expression of glucose metabolism-related factors in fetal liver hepatocytes and regulators of kidney epithelial cell function important for blood pressure regulation. In light of data from our in vivo studies in a guinea pig model of placental insufficiency, we hypothesized that increased trophoblast stress would result in changes to placental-fetal signaling that would influence gene and protein expression in fetal liver and kidney cells. To test this hypothesis, we utilized human cell lines: BeWo choriocarcinoma cells (trophoblast), Human Placental Micro-Vascular Endothelial Cells (HPMVEC; placenta capillary endothelium), and WRL68 (fetal liver hepatocytes) or HEK293T/17 (fetal kidney epithelium), in a multi-layer model designed to mimic cytotrophoblast-villous endothelium-fetal organ communication (Schematic 1).

**Schematic 1.**
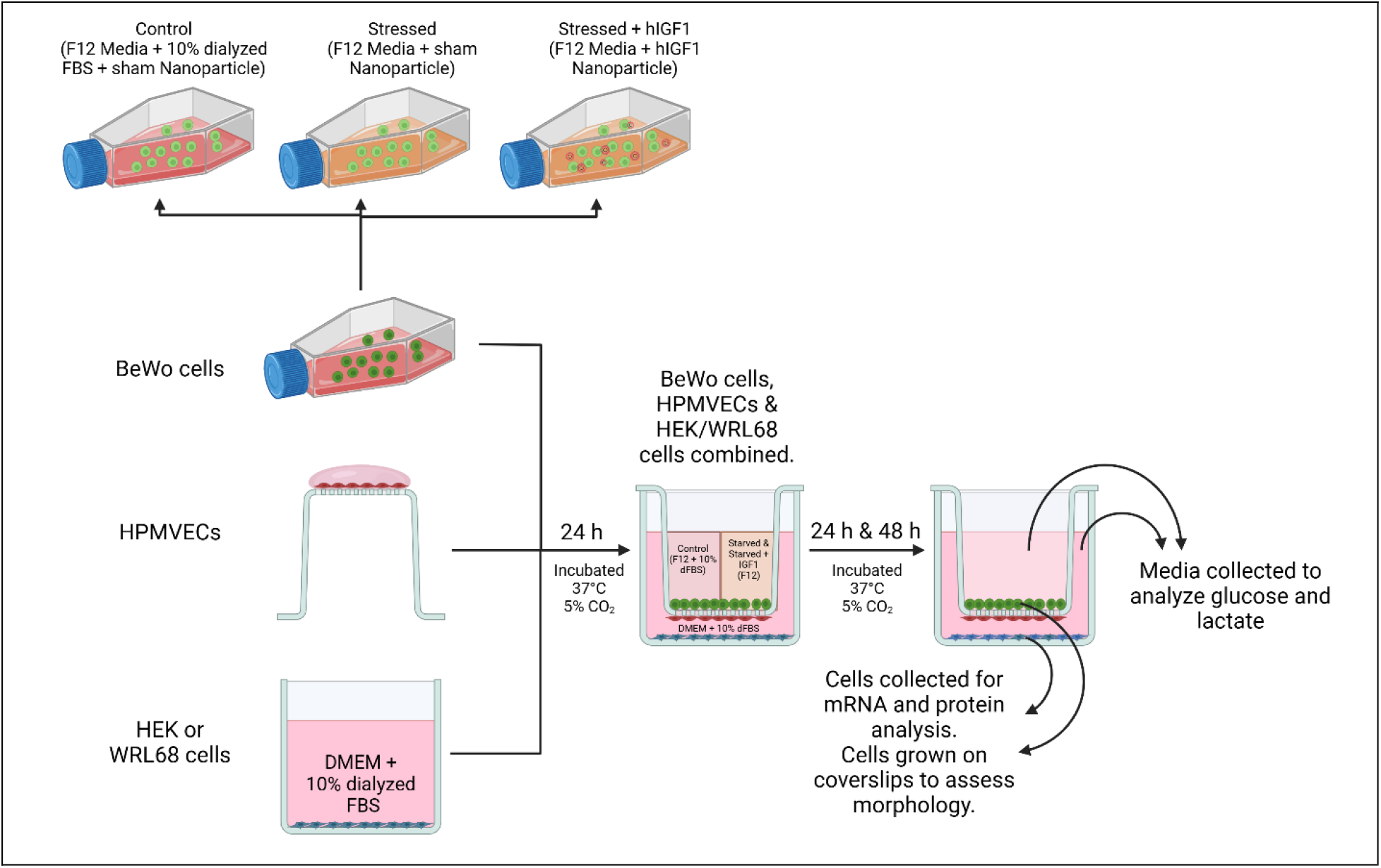
Visual representation of the multi-cell model designed to mimic cytotrophoblast-villous endothelium-fetal organ communication. To alter trophoblast stress response mechanisms prior to co-culture, BeWo cells were treated 1 of 3 ways: Control = growth media with dialyzed FBS + sham nanoparticle; Stressed = growth media without FBS + sham nanoparticle; Stressed+*hIGF1* = growth media without FBS + *human insulin-like 1 growth factor* (*hIGF1*) nanoparticle. After 24 h in treatments (designated 0 h), BeWo cells were placed into the apical transwell chamber of a porous transwell insert. Control BeWo cells were provided media with dialyzed FBS and without nanoparticle. Stressed and Stressed+*hIGF1* BeWo cells were provided media without FBS and without nanoparticle. Human placenta microvascular endothelial cells (HPMVECs) had been grown onto the under-side of the transwell insert for 24 h prior to co-culture. Human fetal liver hepatocytes (WRL68) or human fetal kidney epithelial cells (HEK293T/17) had been placed into the basal chamber of the culture plate for 24 h prior to co-culture. Media in the basal chamber was WRL68 or HEK293T/17 culture media with dialyzed FBS. Cells were co-cultured together for 24 h and 48 h. Culture media from both chambers was collected at 0 h, 24 h and 48 h. BeWo and WRL68 or HEK293T/17 cells were collected at 0 h, 24 h and 48 h to analyzed gene and protein expression.

## Materials and Methods

### Cell Culture Reagents

Gibco’s Ham’s F-12K (Kaighn’s) Medium, Gibco’s DMEM (Dulbecco’s Modified Eagle Medium), Gibco’s heat-inactivated Fetal Bovine Serum (FBS), Gibco’s Trypsin-EDTA (0.05%) and Gibco’s Phosphate Buffered Saline (PBS) were purchased from Thermo Fisher Scientific. Corning’s 0.4 µm, 12 mm, 12-well Transwell Permeable Polycarbonate Membrane Inserts and Plates, Corning’s Penicillin-Streptomycin solution and Cell Application’s Attachment Factor solution were purchased from Fisher Scientific. HyClone™ Dialyzed FBS was purchased from Cytiva. 20 mm sterile, round cover slips were obtained from Quality Biological.

### Nanoparticle Formation

PHPMA_115_-b-PDMEAMA_115_ co-polymer was synthesized and characterized by Dr. Mukesh Gupta (Vanderbilt University)[26]. Plasmids were cloned from a pEGFP-C1 plasmid (Clonetech Laboratories). For the sham nanoparticle, the *CMV* promotor was replaced by a *CYP19A1* promotor and the *GFP* gene was replaced by a non-coding Antisense *GFP* gene. For the *hIGF1* nanoparticle, the *CMV* promotor was replaced by a *CYP19A1* promotor and the *GFP* gene was replaced by a *human IGF1* gene. Lyophilized PHPMA_115_-b-PDMEAMA_115_ co-polymer was reconstituted (10 mg/mL) in sterile saline. Nanoparticles were formed by combining 10 µg of plasmid with 20 µL reconstituted polymer and made to a total volume of 200 µL with sterile saline under aseptic conditions at room temperature.

### Cell Culture

BeWo (CCL98), WRL68 (CL48) and HEK293T/17 (CRL11268) cell lines were purchased from ATCC. Human Placenta Micro-Vascular Endothelial Cells (HPMVEC) were isolated from a normal term placenta as previously reported [33]. BeWo (passages 3-8) cells were maintained in Ham’s F-12 media with 10% FBS and 1% penicillin-streptomycin. WRL68 and HEK293T/17 cells (both passages 3-8) were maintained in DMEM media with 10% FBS and 1% penicillin-streptomycin. HPMVECs (passages 3-8) were maintained on flasks pre-coated with Attachment Factor in DMEM media with 20% FBS and 1% penicillin-streptomycin. All cells were incubated under humidity at 37°C and 5% CO_2_.

### Experimental Procedure

The experimental procedure is outlined in Schematic 1. All experiments were performed under aseptic conditions. On day 1, 1×10^6^ BeWo cells were seeded onto T25 flasks under the following conditions: Control = Ham’s F-12 media + 10% dialyzed FBS (depleted from low-molecular weight components of <10,000 MW) and 1% penicillin-streptomycin, Stressed and Stressed+*hIGF1* = Ham’s F-12 media with 1% penicillin-streptomycin only. Cells were allowed at adhere for 4 hours before sham nanoparticle (Control and Stressed) or *hIGF1* nanoparticle (Stressed+*hIGF1*) was added. 5×10^5^ HPMVECs were seeded onto the under-side of the transwell insert that had been pre-coated with Attachment Factor in 200 µL of normal growth media. Transwell inserts with the HPMVECs were transferred to the incubator with the transwell inserts inverted for 24 h to allow for attachment of the cells. 1×10^5^ WRL68 or HEK293T/17 cells were seeded into the bottom of a 12-well plate and culture in DMEM media with 10% dialyzed FBS and 1% penicillin-streptomycin. On day 2, the culture media covering the HPMVECs was removed and discarded. Culture media from the WRL68 or HEK293T/17 cells was removed, collected and designated timepoint 0 (T0). The HPMVEC transwell inserts were returned to standard culture position within the 12-well plates containing the WRL68 or HEK293T/17 cells. Fresh DMEM media with 10% dialyzed FBS and 1% penicillin-streptomycin was placed into the basolateral chamber of the plate. For the BeWo cells, culture media was removed, collected and designated T0. Cells were washed briefly with PBS to remove any remaining nanoparticles not endocytosed by the cells. BeWo cells were then trypsinized and 1×10^5^ cells were seeded into the inner chamber of the HPMVEC transwell insert. For Control, fresh Ham’s F-12 media with 10% dialyzed FBS and 1% penicillin-streptomycin was added to the apical transwell insert chamber. For Stressed and Stressed+*hIGF1*, fresh Ham’s F-12 media with 1% penicillin-streptomycin only was added to the apical transwell insert chamber. Remaining BeWo cells not seeded into the transwell insert were collected for RNA extraction and confirmation of increased *hIGF1* gene expression. On day 3 (T24) and day 4 (T48), BeWo and WRL68 or HEK293T/17 cells were collected for RNA extraction and protein extraction. WRL68 or HEK293T/17 cells cultured on cover-slips placed in the bottom of the 12-well plate prior to seeding on day 1 were fixed for morphological analysis. Media from both the apical transwell insert chamber (BeWo) and basolateral transwell chamber (HPMVEC with WRL68 or HEK293T/17) was also collected at T24 and T48.

### RNA Extractions and Quantitative PCR (qPCR)

After media was collected from the apical and basolateral transwell insert chambers, HPMVEC cells were removed from the underside of the transwell insert using a clean Kimwipe to prevent potential contamination. BeWo and WRL68 or HEK293T/17 cells were washed was PBS and lysed with 350 µL of RLT buffer (Qiagen). RNA was extracted using the RNeasy Mini Plus kit (Qiagen) following standard manufacturers protocol. A total of 1 µg of RNA was converted to complementary DNA (cDNA) using the High-capacity cDNA Reverse Transcription kit (Applied Biosystems) and diluted to 1:100. For qPCR, 2.5 µl of cDNA was mixed with 10 µl of PowerUp SYBR green (Applied Biosystems), 1.2 µl of primers at a concentration of 10 nM, and water to make up a total reaction volume of 20 µl. Primer sequences are provided in Table 1. Stability of reference genes *ACTB, TBP, HPRT* and *B2M* in the cells was determined using Normfinder [34], and gene expression was normalized to the geometric mean of all four. Reactions were performed using the Quant3 Real-Time PCR System (*Applied Biosystems*), and relative mRNA expression calculated using the comparative CT method with the Design and Analysis Software v2.6.0 (*Applied Biosystems*).

### Western Blot

BeWo and WRL68 or HEK293T/17 cells were prepared for protein extractions as described above for RNA extractions. Cells were lysed in 300 uL of RIPA buffer containing protease and phosphatase inhibitors. Lysates were centrifuged, supernatant collected and protein concentrations determined using Pierce™ Coomassie Plus Assay Kit (*Thermo Fisher Scientific*) following manufacturer’s protocol. 15-20 μg of protein was mixed with Bolt® SDS Loading Buffer (*Invitrogen*) and Reducing Agent (*Invitrogen*) and denatured by heating at 95°C for 10 min. The lysates and a pre-stain protein ladder (PageRuler, *Thermo Fisher Scientific*) were then run on a 4-12% Tris-Bis precast gel (*Invitrogen*) following manufacturers protocols and transferred onto nitrocellulose membranes using the Bolt Mini Electrophoresis unit (*Invitrogen*). Membranes were placed into 5% skim-milk in Tris-buffered Saline containing Tween 20 (TBS-T) and incubated overnight at 4°C. Primary antibodies are provided in Table 1 and were applied for 2 h at room temperature or overnight at 4°C. The membranes were then washed 3 times in fresh TBS-T, and then further incubated with a HRP conjugated secondary (*Cell Signaling 7074* and *7076*, 1:2000) for 2 h at room temperature. Protein bands were visualized by chemiluminescence using SuperSignal West Femto Maximum Sensitivity Substrate (*Thermo Fisher Scientific*) on the Chemidoc Imager (*Bio-Rad*) and signal intensity of the protein bands calculated using Image Lab software (version 6.1, *Bio-Rad*), normalized to bActin. Representative, uncropped blot images are provided in Supplemental Figure S11.

### Media Analysis

Glucose and lactate in BeWo and WRL68 or HEK293T/17 culture media was measured using the YSI model 2700 Glucose/Lactate Analyzer following standard protocols and normalized to total protein.

### Statistics

All experiments were performed in triplicate on 6 independent passages (n = 6 passages). All statistical analyses were performed using SPSS Statistics 29 software. Distribution assumptions were checked with a Q-Q-Plot. Statistical significance was determined using Generalized Linear Modelling with treatment of the BeWo cells (Control, Stressed or Stressed+*hIGF1*) the main effect. Statistical significance was considered at P≤0.05. For statistically significant results, a Bonferroni post hoc analysis was performed. Results are reported as estimated marginal means ± standard error of the mean (SEM).

## Results

### Expression of cell stress markers were elevated in BeWo cells after 24h culture in media without FBS and prevented with IGF1 over-expression

To determine whether culturing BeWo cells in media without FBS and with *hIGF1* nanoparticle treatment resulted changes to BeWo cell function, we analyzed expression of cell stress and DNA damage markers, and glucose metabolism-related factors after 24 h in culture. Increased *IGF1* gene expression after 20 h of *IGF1* nanoparticle treatment was confirmed in the Stressed+*hIGF1* BeWo cells when compared to sham-treated Control and sham-treated Stressed and remained increased throughout the experimental period for BeWo cells co-culture with WRL68 and BeWo cells co-cultured with HEK293T/17 cells (Supplemental Figure S1A & S1B). *IGF2* expression was similar between sham-treated Control, sham-treated Stressed and Stressed+*hIGF1* and did not change across the co-culture period (Supplemental Figure S1C & S1D). In sham-treated Stressed BeWo cells, after 24 h culture in media without FBS, RNA levels of cell stress markers *NFE2L2* and *TP53*, and antioxidants *SOD1* and *SOD2* were increased compared to Control (Figure 1A-D, respectively). Protein expression of DNA damage marker H2A.X(S139) was also increased compared to Control (Figure 1E), whilst expression of Catalase trended towards an increase when compared to Control (Figure 1F).

**Figure 1.**
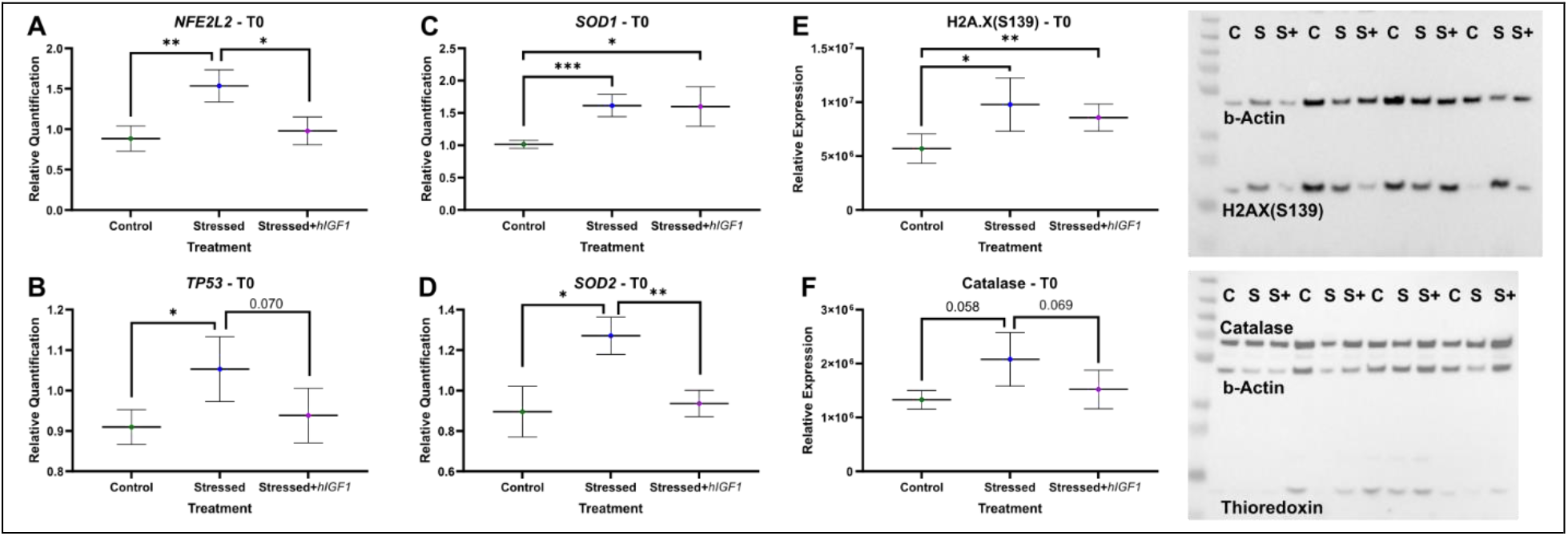
Expression of cellular stress markers in BeWo cells following 24 h culture in stressed (no FBS) media and 20 h of sham or *insulin-like 1 growth factor* (*hIGF1*) nanoparticle treatment. A. Gene expression of cellular defense transcription factor *NFE2L2* was increased in sham treated Stressed BeWo cells when compared sham Control and Stressed+*hIGF1*. **B**. Gene expression of cell stress response transcription factor *TP53* was increased in sham treated Stressed BeWo cells when compared sham Control and trended towards and increase when compared to Stressed+*hIGF1*. **C**. Gene expression of antioxidant *SOD1* was increased in sham Stressed and Stressed+*hIGF1* BeWo cells when compared to sham Control. **D**. Gene expression of antioxidant *SOD2* was increased in sham treated Stressed BeWo cells when compared sham Control and Stressed+*hIGF1*. **E**. Protein expression of DNA damage marker H2A.X(S139) was increased in sham Stressed and Stressed+*hIGF1* BeWo cells when compared to sham Control. Representative Western blot is provided. **F**. Protein expression of antioxidant Catalase trended towards an increase in sham treated Stressed BeWo cells when compared to sham Control and Stressed+*hIGF1*. Data are estimated marginal mean ± SEM calculated using generalized linear modelling. n = 12 independent passages. *P<0.05; **P<0.01; ***P<0.001. NFE2L2: nuclear factor, erythroid 2-like 2. TP53: tumor protein 53. SOD1/2: superoxide dismutase 1/2. H2A.X(S139): phospho-Histone H2A.X. Representative western blot images: C = Control, S = Stressed, S+ = Stressed+*hIGF1*.

In Stressed+*hIGF1* BeWo cells, *NFE2L2* expression was decreased when compared to Stressed but similar to Control (Figure 1A). *TP53* trended towards a decrease when compared to Stressed and was not different to Control (Figure 1B). In contrast, *SOD1* expression reflected Stressed levels and was increased compared to Control (Figure 1C). Whereas *SOD2* expression decreased when compared to Stressed and was similar to Control (Figure 1D). Protein expression of H2A.X(S139) was similar to Stressed cells and increased compared to Control (Figure 1E) whilst Catalase protein expression trended toward a decrease compared to Stressed (Figure 1F). There was no difference in protein expression of antioxidant Thioredoxin with either Starvation or *hIGF1* nanoparticle treatment and protein expression of cleaved-PARP was not detected (data not shown).

### Stressed BeWo cells utilized less glucose from the media and produce more lactate after 24 h in culture, which was not prevented by IGF1 over-expression

Changes in glucose and lactate concentration, normalized to total protein of BeWo cells, was assessed across the 24 h BeWo cells were cultured in stressed (no FBS) and 20 h of *hIGF1* nanoparticle treatment and prior to co-culture. Glucose concentration in Control BeWo media (with FBS: 60.8 mg/dL or 3.4 mM) and Stressed/Stressed+*hIGF1* BeWo media (without FBS: 63.7 mg/dL or 3.5 mM) without the presence of cells was similar. Control BeWo media contained trace amounts of lactate (0.14 mg/dL) whilst Stressed/Stressed+*hIGF1* BeWo media contained no lactate. In Control BeWo cells, glucose concentration in the media decreased 42% and lactate concentration increased to 15.36 mg/dL in 24 h (Table 1). In sham Stressed and Stressed+*hIGF1* BeWo cells, glucose concentration in the media decreased 25% and 23%, respectively, and was higher compared to Control BeWo cell media after 24 h (Table 1). Lactate concentration in the media of Stressed and Stressed+*hIGF1* increased to 19.8 mg/dL and 20.28 mg/dL, respectively, and was higher than levels in the media of Control BeWo cells (Table 1).

In Stressed BeWo cells, gene expression of glucose transporter *SLC2A1* was similar to Control but trended for an increase when compared to Stressed+*hIGF1* (Figure 2A). Gene expression of gluconeogenesis enzyme *G6PC* was increased in Stressed and Stressed+*hIGF1* BeWo cells compared to Control (Figure 2B). *FBP1* and *PCK2* expression were increased in Stressed+*hIGF1* BeWo cells compared to Control and Stressed (Figure 2C and 2D, respectively). *PCK1* mRNA was not detected (no amplification after 40 qPCR cycles) in BeWo cells (data not shown). Gene expression of glycolysis enzymes *HK2, PFKP* and *LDHA* were similar between Control and Stressed BeWo cells but increased in Stressed+*hIGF1* (Figure 2E-G, respectively). Expression of *LDHB* and *LDHC* was increased in Stressed and Stressed+*hIGF1* BeWo cells compared to Control (Figure 2H and 2I, respectively).

**Figure 2.**
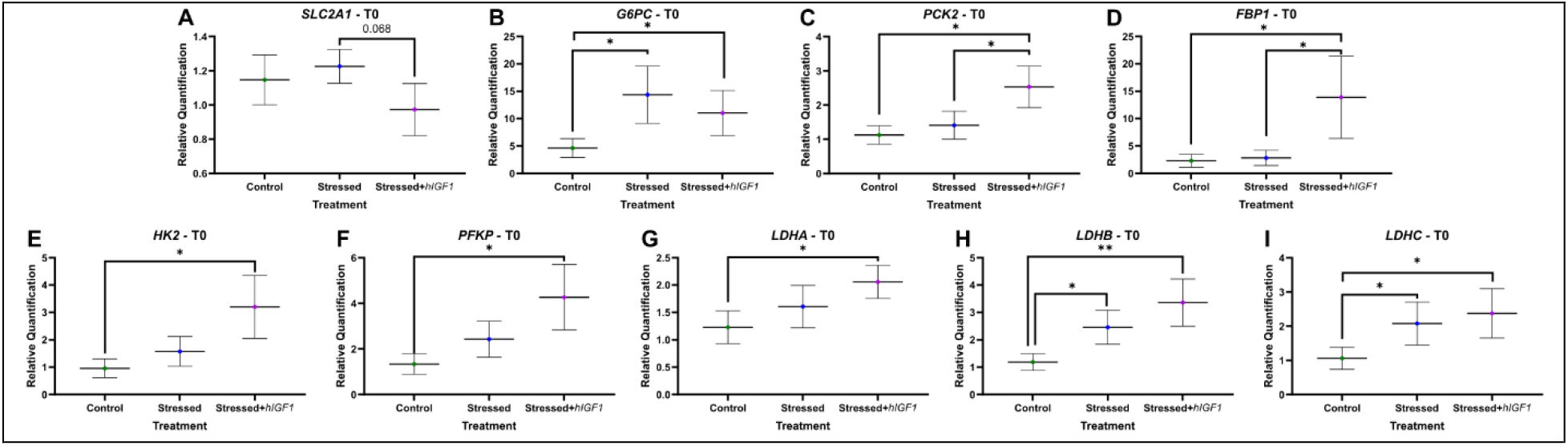
Gene expression of glucose metabolism-related factors in BeWo cells following 24 h culture in stressed (no FBS) media and 20 h of sham or *insulin-like 1 growth factor* (*hIGF1*) nanoparticle treatment. A. Gene expression of glucose transporter *SLC2A1* in Stressed was similar to Control but trended for an increase when compared to Stressed+*hIGF1*. **B**. Gene expression of gluconeogenesis enzyme *G6PC* was increased in Stressed and Stressed+*hIGF1* BeWo cells compared to Control. **C**. *PCK2* expression was increased in Stressed+*hIGF1* BeWo cells compared to Control and Stressed. **D**. *FBP1* expression was increased in Stressed+*hIGF1* BeWo cells compared to Control and Stressed. **E**. Gene expression of glycolysis enzymes *HK2* was similar between Control and Stressed BeWo cells but increased in Stressed+*hIGF1*. **F**. Gene expression of *PFKP* was similar between Control and Stressed BeWo cells but increased in Stressed+*hIGF1*. **G**. Gene expression of *LDHA* was similar between Control and Stressed BeWo cells but increased in Stressed+*hIGF1*. **H**. Expression of *LDHB* was increased in Stressed and Stressed+*hIGF1* BeWo cells compared to Control. **I**. Expression of *LDHC* was increased in Stressed and Stressed+*hIGF1* BeWo cells compared to Control. Data are estimated marginal mean ± SEM calculated using generalized linear modelling. n = 12 independent passages. *P<0.05; **P<0.01; ***P<0.001. SLC2A1: solute carrier family 2 member 1. G6PC: glucose-6-phosphatase. FBP1: Fructose-1,6-bisphosphatase 1. PCK2: Phosphoenolpyruvate carboxykinase 2. HK2: hexokinase 2. PFKP: Phosphofructokinase, platelet. LDHA/B/C: lactate dehydrogenase A/B/C.

### Increasing cell stress mechanisms or over-expressing IGF1 in BeWo cells had no impact on WRL68 gene express of glucose metabolism-related factors

We assessed expression of glucose metabolism-related factors in the WRL68 cell line after 24 h and 48 h in co-culture with BeWo cells and HPMVECs to determine whether altering trophoblast stress responses impacted fetal hepatocytes. There was no difference in the gene expression of *SLC2A1*, gluconeogenesis enzymes (*G6PC, FBP1, FBP2, PCK1, PCK2*), glycolysis enzymes (*HK2, PFKP, LDHA, LDHB, LDHC*) nor cell stress markers (*NFE2L2, TP53*) in WRL68 cells co-culture with Control, Stressed or Stressed+*hIGF1* BeWo cells (Supplemental Figure S2). Irrespective of BeWo treatment, WRL68 gene expression of *SLC2A1, FBP1, FBP2* and *LDHA* was reduced after 24 h in co-culture compared to 0 h but increased back to similar levels as 0 h after 48 h in co-culture (Supplemental Figure S2). Gene expression of *G6PC, FBP2, PCK1, PCK2, HK2, PFKP, LDHB, LDHC, NFE2L2* and *TP53* was reduced in co-cultured WRL68 cells at 24 h compared to 0 h and remained lower at 48 h compared to 0 h (Supplemental Figure S2). Morphologically WRL68 cells expanded and were indistinguishable when comparing co-cultured with Control, Stressed and Stressed+*hIGF1* BeWo cells (Supplemental Figure S3).

### Co-culturing BeWo cells with WRL68 cells was associated with reversal of cell stress responses in BeWo cells and reduced gene expression of gluconeogenesis/glycolysis enzymes

To further understand the implications of co-culturing BeWo, HPMVEC and WRL68 cells, we analyzed expression of cellular stress markers and glucose metabolism-related factors in BeWo cells. In sham-treated Control BeWo cells, gene expression of *NFE2L2* was reduced at 24 h in co-culture and remained lower at 48 h compared to 0 h (Figure 3A). *TP53* expression did not change when comparing 0 h to 24 h but was decreased at 48 h compared to 0 h (Figure 3B). *SOD1* and *SOD2* gene expression were higher at 24 h and 48 h when compared to 0 h (Figure 3C & 3D). Protein expression of H2A.X(S139) remained unchanged across the co-culture with WRL68 cells (Figure 3E), whilst Catalase protein expression decreased at 24 h and 48 h in co-culture when compared to 0 h (Figure 3F).

**Figure 3.**
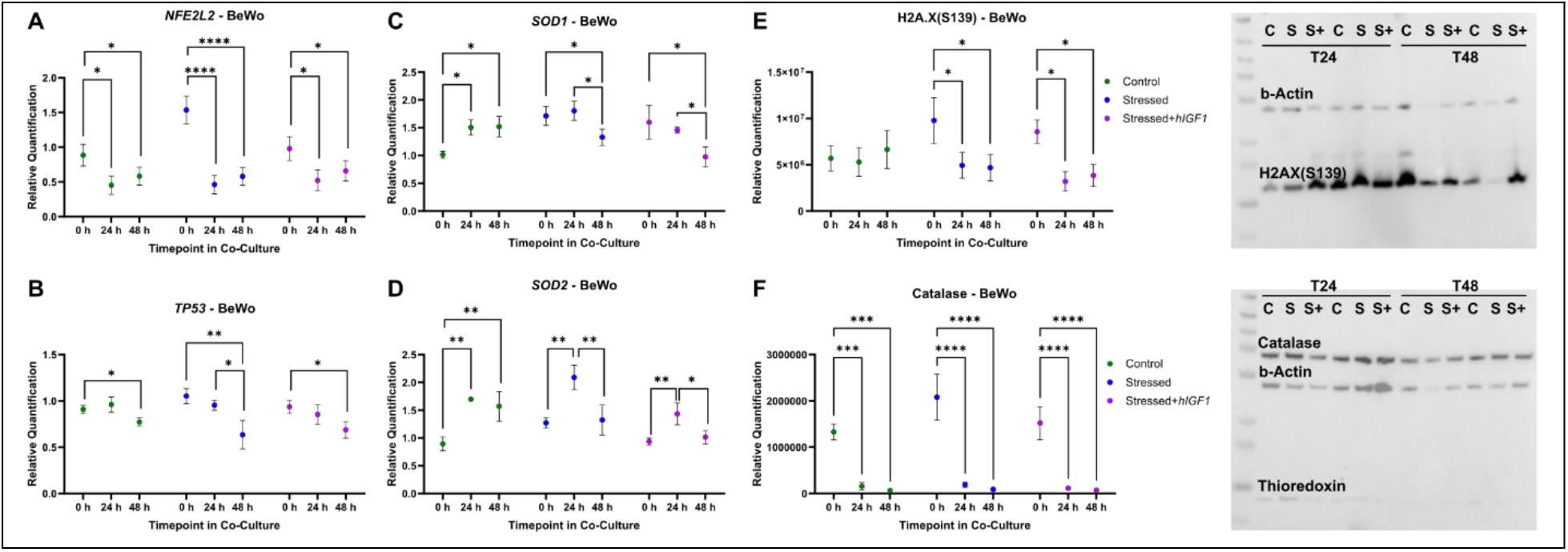
Gene expression of cellular stress markers sham-treated Control BeWo Cells (complete culture media), sham-treated Stressed BeWo cells (no FBS in culture media) or Stressed + *insulin-like 1 growth factor* (*hIGF1*) nanoparticle treated BeWo cells co-cultured with WRL68 cells and HPMVECs. A. Irrespective of BeWo treatment, gene expression of *NFE2L2* was reduced at 24 h in co-culture and remained lower at 48 h compared to 0 h. **B**. For all BeWo treatments, *TP53* expression did not change when comparing 0 h to 24 h but was decreased at 48 h compared to 0 h. **C**. *SOD1* in Control BeWo cells was higher at 24 h and 48 h when compared to 0 h. In Stressed and Stressed+*hIGF1* BeWo cells, *SOD1* expression was lower at 48 h compared to 0 h and 24 h. **D**. *SOD2* in Control BeWo cells was higher at 24 h and 48 h when compared to 0 h. In Stressed and Stressed+*hIGF1* BeWo cells, *SOD2* expression was lower at 48 h compared to 0 h and 24 h. **E**. In Control BeWo cells, protein expression of H2A.X(S139) remained unchanged across the co-culture with WRL68 cells. H2A.X(S139) expression in Stressed and Stressed+*hIGF1* BeWo cells was lower at 24 h and 48 h compared to 0 h. Representative western blot image to the right of graph. **F**. Catalase protein expression decreased at 24 h and 48 h in co-culture when compared to 0 h, irrespective of BeWo treatment. Representative western blot image to the right of graph. Data are estimated marginal mean ± SEM calculated using generalized linear modelling. n = 6 independent passages. *P<0.05; **P<0.01; ***P<0.001. HPMVEC: human placenta microvascular endothelial cells. NFE2L2: nuclear factor, erythroid 2-like 2. TP53: tumor protein 53. SOD1/2: superoxide dismutase 1/2. H2A.X(S139): phospho-Histone H2A.X. Representative western blot images: C = Control, S = Stressed, S+ = Stressed+*hIGF1*.

In co-cultured Stressed BeWo cells, *NFE2L2* was reduced at 24 h and 48 h compared to 0 h, and similar to levels in co-cultured Control BeWo cells at the same timepoints (Figure 3A). *TP53* and *SOD1* expression was lower at 48 h when compared to 0 h and 24 h, and comparable to levels in co-cultured Control BeWo cells at 48 h (Figure 3B & 3C). *SOD2* gene expression was higher at 24 h when compared to 0 h but reduced between 24 h and 48 h (Figure 3D). H2A.X(S139) and Catalase protein expression in Stressed BeWo cells was lower at 24 h in co-culture with WRL68 cells compared to 0 h, remained lower at 48 h, and was comparable to levels in co-culture Control BeWo cells at 48 h (Figure 3E & 3F).

In co-cultured Stressed+*hIGF1* BeWo cells, gene expression of *NFE2L2, TP53* and *SOD2* and protein expression of H2A.X(S139) and Catalase at 24 h and 48 h changed in a similar manner to Stressed BeWo cells (Figure 3). *SOD1* gene expression was lower in co-cultured Stressed+*hIGF1* BeWo cells at 48 h compared to 0 h, and lower than Control BeWo cells at the 48 h timepoint (Figure 3C).

Gene expression of *SLC2A1, G6PC, FBP1*, and *PCK2* in Control BeWo cells did not change when co-cultured with WRL68 cells (Supplemental Figure S4). Gene expression of *HK2* and *PFKP* was increased in Control BeWo cells at 24 h in co-culture with WRL68 cells compared to 0 h and remained higher at 48 h (Supplemental Figure S4). *LDHA* and *LDHC* were lower in co-cultured Control BeWo cells at 48 h compared to 0 h (Supplemental Figure S4). In co-cultured Stressed and Stressed+*hIGF1* BeWo cells, gene expression of gluconeogenesis enzymes that were higher at 0 h were reduced by 48 h, and comparable to Control BeWo cells at 48 h (Supplemental Figure S4). Changes in gene expression of glycolysis enzymes in co-cultured Stressed and Stressed+*hIGF1* BeWo cells occurred in a similar manner across the co-culture period with WRL68 cells and were no different from Control BeWo cells at 48 h (Supplemental Figure S4).

### Co-culture of BeWo and WRL68 cells resulted in increased media glucose concentrations in both the apical and basal transwell insert chambers

In Control BeWo cells co-cultured with WRL68 cells, glucose concentrations were ∼5X higher in the apical transwell chamber (BeWo side) and 30% higher in the basal chamber (WRL68 side) at 24 h when compared to 0 h (Table 1). Between 24 h and 48 h, glucose concentrations reduced by 21% in the apical transwell chamber and 44% in the basal chamber (Table 1). Lactate concentrations increased ∼32,600% in the apical transwell chamber and ∼14,400% in the basal chamber between 0 h and 24 h (Table 1). Lactate concentration increased a further 86% and 69% in the apical transwell chamber and basal chamber, respectively, from 24 h to 48 h (Table 1).

In co-cultured Stressed BeWo cells, glucose concentrations were ∼5.5X higher in the apical transwell chamber and 42% higher in the basal chamber when compared to 0 h, and higher when compared to concentrations in the co-cultured Control chambers at 24 h (Table 1). Glucose concentration in both chambers decreased 22% (apical) and 46% (basal) between 24 h and 48 h, and remained higher at 48 h when compared to co-cultured Control (Table 1). Lactate concentrations went from 0 mg/dL at 0 h to 55 mg/dL at 24 h in the apical transwell chamber, and was higher when compared to concentrations in the Control BeWo apical transwell chamber at 24 h (Table 1). Lactate concentrations in the basal chamber increased ∼13,800% between 0 h and 24 h, and was similar to lactate concentrations in the basal chamber of co-cultured Control (Table 1). Between 24 h and 48 h, lactate concentrations increased 52% in the apical transwell chamber and 70% in the basal chamber, and were similar to concentrations of co-cultured Control at 48 h (Table 1).

In co-cultured Stressed+*hIGF1* BeWo cells, glucose increased ∼5.3X concentrations in the apical transwell chamber between 0 h and 24 h, and was similar to concentrations in the apical transwell chamber of co-cultured Stressed (Table 1). Glucose concentrations in the basal chamber increased 28% from 0 h to 24 h, and were lower than concentrations in the co-cultured Stressed BeWo basal chamber (Table 1). Between 24 h and 48 h, glucose concentrations in both chambers of the co-cultured Stressed+*hIGF1* group decreased in a similar manner as Control and Stressed (Table 1).

### Increasing cell stress mechanisms or over-expressing IGF1 in BeWo cells had no impact on gene expression of angiogenic factors in fetal kidney epithelial HEK293T/17 cells

Similar to further understanding the implications of manipulating trophoblast function on fetal hepatocytes, we assessed the expression of important kidney epithelial cell function factors in the HEK293T/17 cell line after 24 h and 48 h in co-culture with BeWo cells. There was no difference in the gene expression of *NFE2L2, TP53, SOD1, SOD2, TGFb, VEGF, IGF1R* and *ACE* in HEK293T/17 cells co-culture with Control, Stressed or Stressed+*hIGF1* BeWo cells (Supplemental Figure S5). Irrespective of BeWo manipulation, HEK293T/17 gene expression of *NFE2L2, TP53, SOD1, SOD2, TGFb, VEGF* and *IGF1R* was reduced after 24 h in co-culture when compared to 0 h (Supplemental Figure S5). Gene expression of *NFE2L2, TP53, SOD1, SOD2, TGFb* was increased at 48 h in co-culture when compared to 24 h. *VEGF, IGF1R* and *ACE* gene expression was higher at 48 h in co-culture with BeWo cells when compared to 0 h and 24 h. Morphologically HEK293T/17 cells expanded and were indistinguishable when comparing those co-cultured with Control, Stressed and Stressed+*hIGF1* BeWo cells (Supplemental Figure S6).

### Co-culturing BeWo cells with HEK293T/17 cells increased cell stress responses in BeWo cells and gene expression of gluconeogenesis/glycolysis enzymes

To further elucidate whether signals from fetal cells or media glucose concentrations were more influential on changes to trophoblast gene and protein expression, we assessed expression of cellular stress markers and glucose metabolism-related factors in BeWo cells co-cultured with HEK293T/17 cells. In co-cultured Control BeWo cells, gene expression of *NFE2L2* and *TP53* was increased at 24 h and remained higher at 48 h when compared to 0 h (Figure 4A & 4B). *SOD1* and *SOD2* gene expression and Catalase protein expression did not change across the co-culture period with HEK293T/17 cells (Figure 4C, 4D & 4F). Protein expression of H2A.X(S139) increased in at 24 h when compared to 0 h and was still higher at 48 h (Figure 4D).

**Figure 4.**
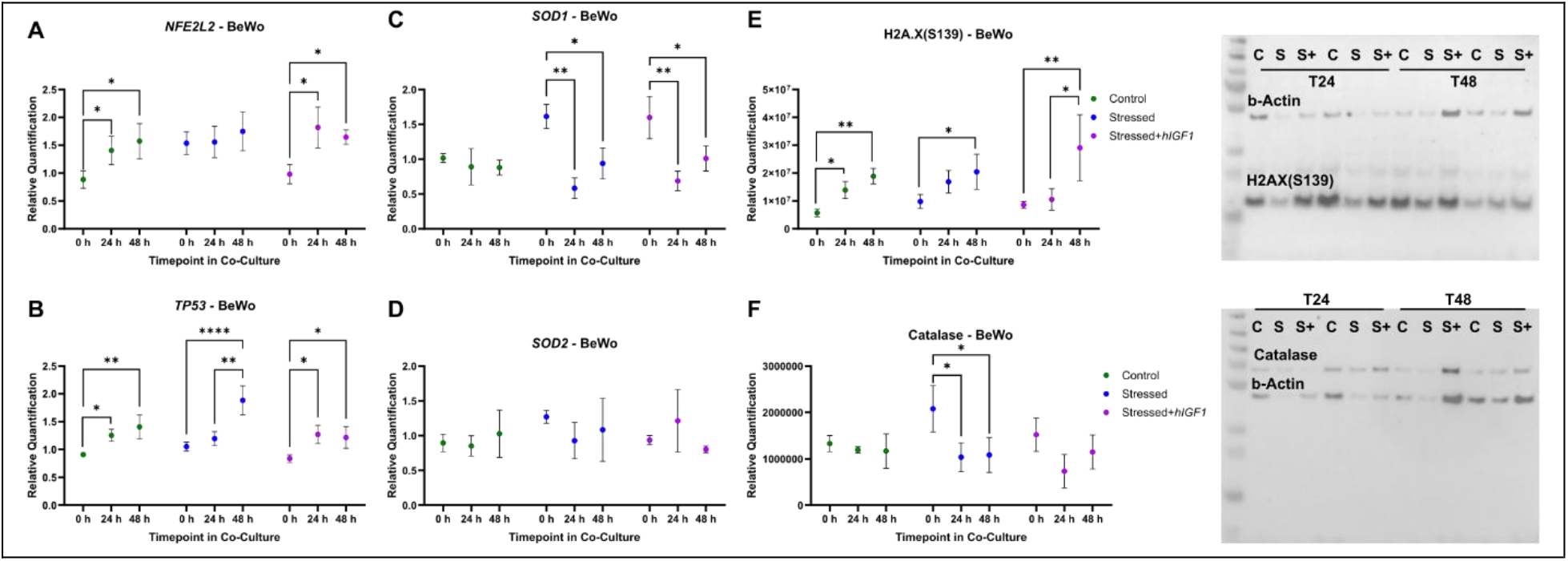
Gene expression of cellular stress markers in sham-treated Control BeWo Cells (complete culture media), sham-treated Stressed BeWo cells (no FBS in culture media) or Stressed + *insulin-like 1 growth factor* (*hIGF1*) nanoparticle treated BeWo cells co-cultured with HEK293T/17 cells and HPMVECs. A. In Control and Stressed+*hIGF1* BeWo cells, gene expression of *NFE2L2* was increased at 24 h and remained higher at 48 h when compared to 0 h. *NFE2L2* expression did not change in Stressed BeWo cells across the co-culture. **B**. In Control and Stressed+*hIGF1* BeWo cells, gene expression of *TP53* was increased at 24 h and remained higher at 48 h when compared to 0 h. In Stressed BeWo cells, expression of *TP53* was higher at 48 h compared to 0 h and 24 h. **C**. *SOD1* gene expression did not change in Control BeWo cells across the co-culture period. *SOD1* expression was decreased at 24 h and 48 h compared to 0 h in Stressed and Stress+*hIGF1* co-cultured BeWo cells. **D**. *SOD2* gene expression did not change across the co-culture period for all BeWo treatments. **E**. Protein expression of H2A.X(S139) increased in at 48 h when compared to 0 h for all BeWo treatments. Representative western blot image to the right of graph. **F**. Catalase protein expression did not change across the co-culture period in Control and Stressed+*hIGF1* BeWo cells. Catalase expression was reduced in the Stressed BeWo cells at 24 h and 48 h compared to 0 h. Representative western blot image to the right of graph. Data are estimated marginal mean ± SEM calculated using generalized linear modelling. n = 6 independent passages. *P<0.05; **P<0.01; ***P<0.001. HPMVEC: human placenta microvascular endothelial cells. NFE2L2: nuclear factor, erythroid 2-like 2. TP53: tumor protein 53. SOD1/2: superoxide dismutase 1/2. H2A.X(S139): phospho-Histone H2A.X. Representative western blot images: C = Control, S = Stressed, S+ = Stressed+*hIGF1*.

In Stressed BeWo cells, *NFE2L2* did not change across the co-culture period with HEK293T/17 cells and was comparable to elevated levels in co-cultured Control BeWo cells (Figure 4A). *TP53* was higher in Stressed BeWo cells at 48 h in co-culture when compared to 0 h and 24 h, and higher than sham Control BeWo cells at 48 h (Figure 4B). *SOD1* gene expression was lower in sham Stressed BeWo cells at 24 h and 48 h in co-culture with HEK293T/17 cells compared to 0 h, and similar when compared to sham Control BeWo cells at the same time point (Figure 4C). *SOD2* gene expression did not change across the co-culture period (Figure 4D). H2A.X(S139) protein expression was higher in sham Stressed BeWo cells at 48 h in co-culture with HEK293T/17 cells whilst Catalase protein expression was lower at 24 h in co-culture with HEK293T/17 cells compared to 0 h, and remained lower at 48 h (Figure 4E & 4F).

In Stressed+*hIGF1* BeWo cells, gene expression of *NFE2L2* and *TP53* at 24 h and 48 h in co-culture with HEK293T/17 cells changed in a similar manner to sham Control BeWo cells (Figure 4A & 4B). *SOD1* gene expression was lower in Stressed+*hIGF1* BeWo cells at 24 h and 48 h in co-culture with HEK293T/17 cells compared to 0 h, and similar when compared to sham Control and sham Stressed BeWo cells (Figure 4C). *SOD2* gene expression and Catalase protein expression did not change in Stressed+*hIGF1* BeWo cells across the culture period with HEK293T/17 (Figure 4D & 4F). Protein expression of H2A.X(S139) was higher in Stressed+*hIGF1* BeWo cells at 48 h when compared to 0 h and 24 h and similar to levels in sham Control and sham Stressed at 48 h (Figure 4E).

Gene expression of *SLC2A1* was increased in BeWo cells co-cultured with HEK293T/17 cells at 24 h, and further again at 48 h when compared to 0 h, irrespective of BeWo treatment (Figure 5A). Gene expression of *G6PC, FBP1, PCK2, HK2, PFKP, LDHA, LDHB*, and *LDHC* were increased by 48 h in sham Control BeWo cells co-cultured with HEK293T/17 when compared to 0 h (Figure 5B-5I). In sham Stressed and Stressed+*hIGF1* BeWo cells co-cultured with HEK293T/17 cells, *G6CP* gene expression did not change at 24 h and 48 h, and were comparable to elevated expression levels in sham Control BeWo cells (Figure 5B). Gene expression of *FBP1* and *PCK2* were higher in sham Stressed BeWo cells by 48 h in co-culture with HEK293T/17 cells when compared to 0 h, but unchanged in Stressed+*hIGF1* at 24 h and 48 h compared to 0 h (Figure 5C & 5D).

**Figure 5.**
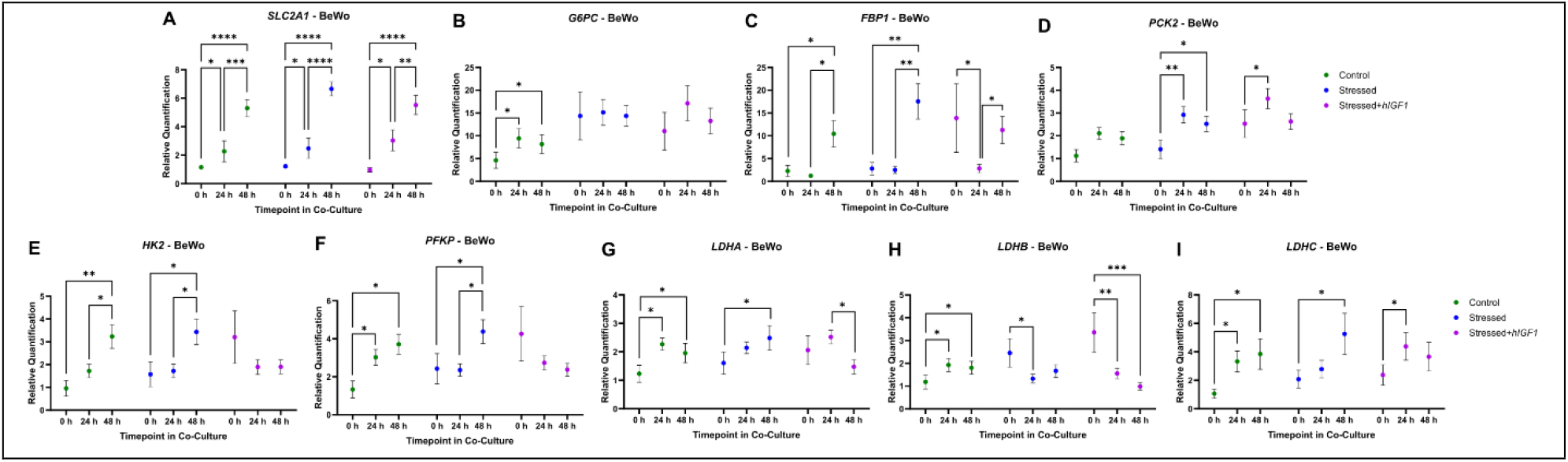
Gene expression of glucose metabolism-related factors in sham-treated Control BeWo Cells (complete culture media), sham-treated Stressed BeWo cells (no FBS in culture media) or Stressed + *insulin-like 1 growth factor* (*hIGF1*) nanoparticle treated BeWo cells co-cultured with HEK293T/17 cells and HPMVECs. A. Gene expression of *SLC2A1* was increased in BeWo cells co-cultured with HEK293T/17 cells at 24 h, and further again at 48 h when compared to 0 h, irrespective of BeWo treatment. **B**. In Control BeWo cells, gene expression of *G6PC* increased at 24 h when compared to 0 h and remained higher at 48 h compared to 0 h. In sham Stressed and Stressed+*hIGF1* BeWo cells, *G6CP* gene expression did not change at 24 h and 48 h compared to 0 h. **C**. In Control and Stressed BeWo cells, gene expression of *FBP1* increased at 48 when compared to 0 h and 24 h. *FBP1* expression was lower in Stressed+*hIGF1* BeWo cells at 24 h compared to 0 h and 48 h. **D**. In Control and Stressed BeWo cells, *PCK2* increased at 24 h compared to 0 h, and remained increased at 48 h compared to 0 h. *PCK2* expression was higher in Stressed+*hIGF1* BeWo cells at 24 h compared to 0 h and 48 h. **E**. In Control and Stressed BeWo cells, *HK2* increased at 48 h when compared to 0 h and 24 h. **F**. In Control BeWo cells, *PFPK* increased at 48 h when compared to 0 h and 24 h. In Stressed BeWo cells, *PFKP* expression increased at 48 h compared to 0 h and 24 h. Expression of *PFKP* in Stressed+*hIGF1* BeWo cells remained unchanged across the co-culture period. **G**. In Control BeWo cells, *LDHA* increased at 24 h when compared to 0 h, and remained increased at 48 h compared to 0 h. In Stressed BeWo cells, *LDHA* increased at 48 h compared to 0 h. In Stressed+*hIGF1* BeWo cells, *LDHA* decreased at 48 h compared to 24 h. **H**. In Control BeWo cells, *LDHB* increased at 24 h when compared to 0 h, and remained increased at 48 h compared to 0 h. In Stressed BeWo cells, *LDHB* decreased at 24 h compared to 0 h. In Stressed+*hIGF1* BeWo cells, *LDHB* decreased at 24 h and 48 h compared to 0 h. **I**. In Control BeWo cells, *LDHC* increased at 24 h when compared to 0 h, and remained increased at 48 h compared to 0 h. In Stressed BeWo cells, *LDHC* increased at 48 h compared to 0 h. In Stressed+*hIGF1* BeWo cells, *LDHC* increased at 24 h compared to 0 h. Data are estimated marginal mean ± SEM calculated using generalized linear modelling. n = 6 independent passages. *P<0.05; **P<0.01; ***P<0.001. HPMVEC: human placenta microvascular endothelial cells. SLC2A1: solute carrier family 2 member 1. G6PC: glucose-6-phosphatase. FBP1: Fructose-1,6-bisphosphatase 1. PCK2: Phosphoenolpyruvate carboxykinase 2. HK2: hexokinase 2. PFKP: Phosphofructokinase, platelet. LDHA/B/C: lactate dehydrogenase A/B/C.

### Co-culture of BeWo and HEK293T/17 cells resulted in increased media glucose concentrations in the apical insert chamber but not the basal chamber

In sham Control BeWo cells co-cultured with HEK293T/17, glucose concentrations in the apical transwell chamber (BeWo side) were ∼4.5X higher but only 4% higher in the basal chamber (HEK293T/17) at 24 h when compared to 0 h (Table 1). Glucose concentrations in the apical transwell chamber reduced by 12% and 42% in the basal chamber between 24 h and 48 h (Table 1). Lactate concentrations in the apical transwell chamber of sham Control BeWo cells increased ∼22,900% and ∼8,300% in the basal chamber between 0 h and 24 h (Table 1). Lactate concentration increased a further 92% and 60% in the apical transwell chamber and basal chamber, respectively, from 24 h to 48 h (Table 1).

In sham Stressed BeWo cells, glucose concentrations in the apical transwell chamber were ∼3.6X higher and did not change in the basal chamber after 24 h when compared to 0 h, and were lower when compared to sham Control at 24 h (Table 1). Glucose concentration in both chambers decreased 6% (apical) and 44% (basal) between 24 h and 48 h, and were similar at 48 h when compared to sham Control (Table 1). Lactate concentrations in the apical transwell chamber of sham Stressed BeWo cells went from 0 mg/dL at 0 h to 25 mg/dL at 24 h, and was lower when compared to sham Control BeWo cells at 24 h (Table 1). Lactate concentrations in the basal chamber of sham Stressed BeWo cells increased ∼7,950% between 0 h and 24 h, and was similar to lactate concentrations in the basal chamber of sham Control (Table 1). Between 24 h and 48 h, lactate concentrations in the apical transwell chamber of sham Stressed BeWo cells increased 101% but was still lower than in the apical transwell chamber of sham Control BeWo cells (Table 1). In the basal chamber, lactate concentration increased by 70%, and was similar to levels in the basal chamber of sham Control at 48 h (Table 1).

In Stressed+*hIGF1* BeWo cells, glucose concentrations in the apical transwell chamber increased ∼5X between 0 h and 24 h, and was higher than concentrations in the apical transwell chamber of sham Control and sham Stressed BeWo cells at 24 h (Table 1). Glucose concentrations in the basal chamber increased 6% from 0 h to 24 h, and were similar to sham Control and sham Stressed BeWo basal chamber at 24 h (Table 1). Between 24 h and 48 h, glucose concentrations in both chambers of the Stressed+*hIGF1* BeWo cells decreased in a manner and were no different from sham Control or sham Stressed, and concentrations in both chambers was similar to sham Control and sham Stressed at 48 h (Table 1).

## Discussion

Fetal growth requires multi-directional maternal-placental-fetal communication, and dysfunction of any aspect of this communication can result in perturbation of fetal development with life-long consequences. In this proof-of-concept study, we aimed to assess the impacts of altered trophoblast stress response mechanisms and *hIGF1* nanoparticle treatment on fetal liver hepatocytes and fetal kidney epithelial cells. We confirmed that after 24 h culture in media without FBS, expression levels of trophoblast cellular stress mechanisms were elevated as well as expression of gluconeogenesis rate-limiting enzymes. When trophoblast cells were stressed but *IGF1* over-expressed with a transient *hIGF1* nanoparticle gene therapy, cellular stress mechanisms were not increased, however upregulation of enzymes in the gluconeogenesis and glycolysis pathways remained. Our original hypothesis was that altering trophoblast stress mechanisms would result in changes to placental-fetal signaling that would influence gene and protein expression in fetal liver and kidney cells; this was not observed. Unexpectedly, we found a potential for trophoblast to not only transport glucose but also produce glucose, which goes beyond the scope of the current manuscript but worth further investigation. Overall, this study furthers our understanding of the mechanisms linking placental function and fetal glucose status in fetal developmental programming and shaping long-term health outcomes.

Placental insufficiency is often associated with dysfunctional trophoblast phenotypes [35]. Increased oxidative stress and defects in trophoblast proliferation and differentiation pathways are pathogenic hallmarks. In the current study, we confirmed that culturing BeWo cells for 24 h in media without the support of FBS resulted in increased expression of cell stress and DNA damage markers as well as reduced utilization of glucose from the media. Increased expression of gluconeogenesis and glycolysis enzyme genes in the trophoblast was also observed which has also been shown in term placentas and associated with mitochondrial dysfunction [36, 37]. We speculate that upregulation in the gene expression of these gluconeogenesis/glycolysis enzymes is a cell response to redirect resources from the TCA cycle to other processes, however further investigations beyond the scope of the current manuscript are required. Increased expression of *IGF1* with the *hIGF1* gene therapy increased expression of additional enzymes within the gluconeogenesis and glycolysis pathways whilst also preventing increased expression of cell stress markers. We have previously shown that *hIGF1* gene therapy protected against increased cell death and decreased mitochondrial activity in BeWo cells stressed with hydrogen peroxide [29]. Altogether further confirming the positive effect of increased *IGF1* expression on molecular mechanisms in trophoblast cells.

The original aim of this study was to determine how increased trophoblast stress would result in changes to placental-fetal signaling that would influence gene and protein expression in fetal liver and kidney cells. In our BeWo-HPMVEC-fetal liver WRL68 co-culture model we identified a significant increase in glucose within the apical transwell BeWo culture media within 24 h of co-culture which was higher when Stressed BeWo cells were present. This was likely due to the transport of glucose, facilitated by specific glucose transporters like SLC2A1/GLUT1 in the BeWo down a concentration gradient from the high glucose media in the basal compartment [38]. In addition, liver hepatocytes can produce glucose via gluconeogenesis [39]. Whilst we can only speculate that the increased glucose within the co-culture media at 24 h was due to WRL68 production, we hypothesize that there is signaling from the BeWo cells that rapidly increases glucose production within the WRL68 cells resulting in alleviation of cell stress mechanisms within the BeWo cells. We chose not to reduce glucose in the WRL68 culture media as our aim was to specifically look at manipulation of the trophoblast. However, whether similar outcomes would be observed if glucose was also lower in the basal chamber of the co-culture system is an interesting future direction.

Co-culture of BeWo, HPMVECs and fetal kidney epithelial HEK293T/17 cells showed no difference in HEK293T/17 morphology or expression of key genes relating to angiogenesis and blood pressure regulation with BeWo manipulation. However, whilst co-cultured BeWo and WRL68 cells showed decreased cellular stress mechanisms in stressed BeWo cells, cellular stress mechanisms were increased in Control BeWo cells when co-cultured with HEK293T/17. Additionally, there was a significant increase in glucose within the media of the apical transwell containing the BeWos within 24 h of co-culture that was associated with increased gene expression of glycolysis and gluconeogenesis enzymes in BeWo cells by 48 h. Upregulation of glycolysis and gluconeogenesis enzymes expression was not observed when BeWo cells were co-cultured with WRL68 cells and thus provides evidence for the possibility that trophoblast cells, in addition to transporting glucose, can produce glucose. The idea that trophoblast can produce glucose in addition to transporting it is not novel but often debated [40]. Efforts to determine whether the placenta has the capacity to produce glucose have focused on G6PC, the key enzyme which catalyzes the terminal reaction of glucose from glucose-6-phosphate [41]. Whilst most highly expressed and active in the liver, G6PC activity is not liver exclusive and has been found in other tissues including the placenta [42]. It is also possible that in this study, increased glucose in the apical transwell media is from BeWo transport of glucose from stores within the HPMVECs and/or HEK293T/17 cells and requires further investigation beyond the current scope.

One of the strengths of this study was the utilization of a culture system which mimics the cytotrophoblast-endothelial environment. The structure of the chorionic villous tissue is complex, requiring multiple cell types with highly specialized functions to work cooperatively to coordinate nutrient, oxygen and waste exchange between maternal and fetal circulations. Nutrients from the maternal blood and trophoblast-derived endocrine factors must cross a layer of endothelial cells prior to entering the placental-fetal circulation, hence we chose to include HPMVECs within the culture system. However, our hypothesis centered on the impact of stressing or stressing and treating trophoblast with a *hIGF1* gene therapy on fetal liver and kidney cells, with the HPMVECs only present as an intermediary cell type to better recapitulate the in vivo situation. As we did not collect or analyze the HPMVECs, we cannot be certain of their influence on the outcomes observed in both the BeWo and WRL68 or HEK293T/17 cells. As this was a proof-of-concept study, further investigations into the absolute role of the HPMVECs or whether outcomes would be different if syncytiotrophoblast were also incorporated are beyond the scope but present interesting future directions.

In conclusion, we hypothesized that increased trophoblast stress would result in changes to placental-fetal signaling that would influence gene and protein expression in fetal liver and kidney cells. Our data did not support this hypothesis but instead demonstrated that cytotrophoblast under stress turn on mechanisms involved in glucose production. Whether this is reflected in vivo remains uninvestigated but may represent a placental compensation mechanism in complicated pregnancies. Most importantly, this study highlights the complexities of the placenta and the often-overlooked challenges in studying such a dynamic and multifaceted system.

## Acknowledgments

We would like to thank Aly Williams for her experimental assistance.

## Author Contributions

HNJ: Conceptualization, Resources, Funding Acquisition, Writing – Review & Editing. RLW: Conceptualization, Methodology, Validation, Investigation, Formal Analysis, Writing - Original Draft, Project Administration, Funding Acquisition

