## Supplemental Material for "A human cytotrophoblast-villous endothelium-fetal organ multi-cell model and the impact on gene and protein expression in placenta cytotrophoblast, fetal hepatocytes and fetal kidney epithelial cells"

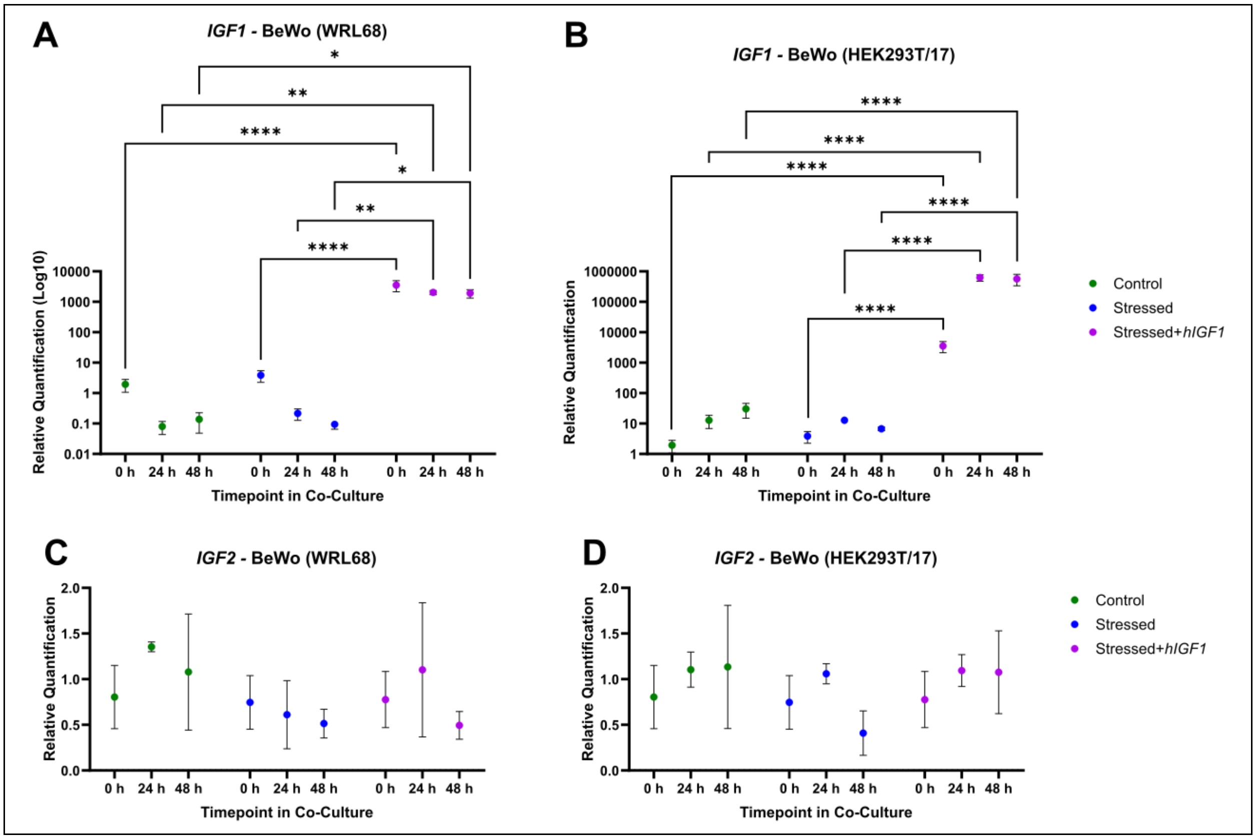


**Supplemental Figure S1. *Insulin-like Growth Factor 1* (*IGF1*) and *IGF2* gene expression in sham-treated Control BeWo Cells (complete culture media), sham-treated Stressed BeWo cells (no FBS in culture media) or Stressed + *insulin-like 1 growth factor* (*hIGF1*) nanoparticle treated BeWo cells co-cultured with WRL68 cells and HPMVECs. A & B.** At 0 h and prior to cell culture with WRL68 or HEK293T/17 cells, *hIGF1* nanoparticle treatment increased *IGF1* expression in Stressed+*hIGF1* BeWo cells after 20 h in culture when compared to sham treated Control and sham treated Stressed BeWo cells. Even after a complete media change to remove any remaining *hIGF1* nanoparticle in the culture media, *IGF1* expression remained elevated in BeWo cells co-cultured with WRL68 or HEK293T/17 cells at 24 h and 48 h. **C & D**. *IGF2* gene expression in BeWo cells was no different between sham treated Control, sham treated Stressed or Stressed+*hIGF1* treatment across the culture period. Data are estimated marginal mean ± SEM calculated using generalized linear modelling. n = 12 independent passages (0 h) and n = 6 independent passages (24 h and 48 h). *P<0.05; **P<0.01; ***P<0.001; ****P<0.0001

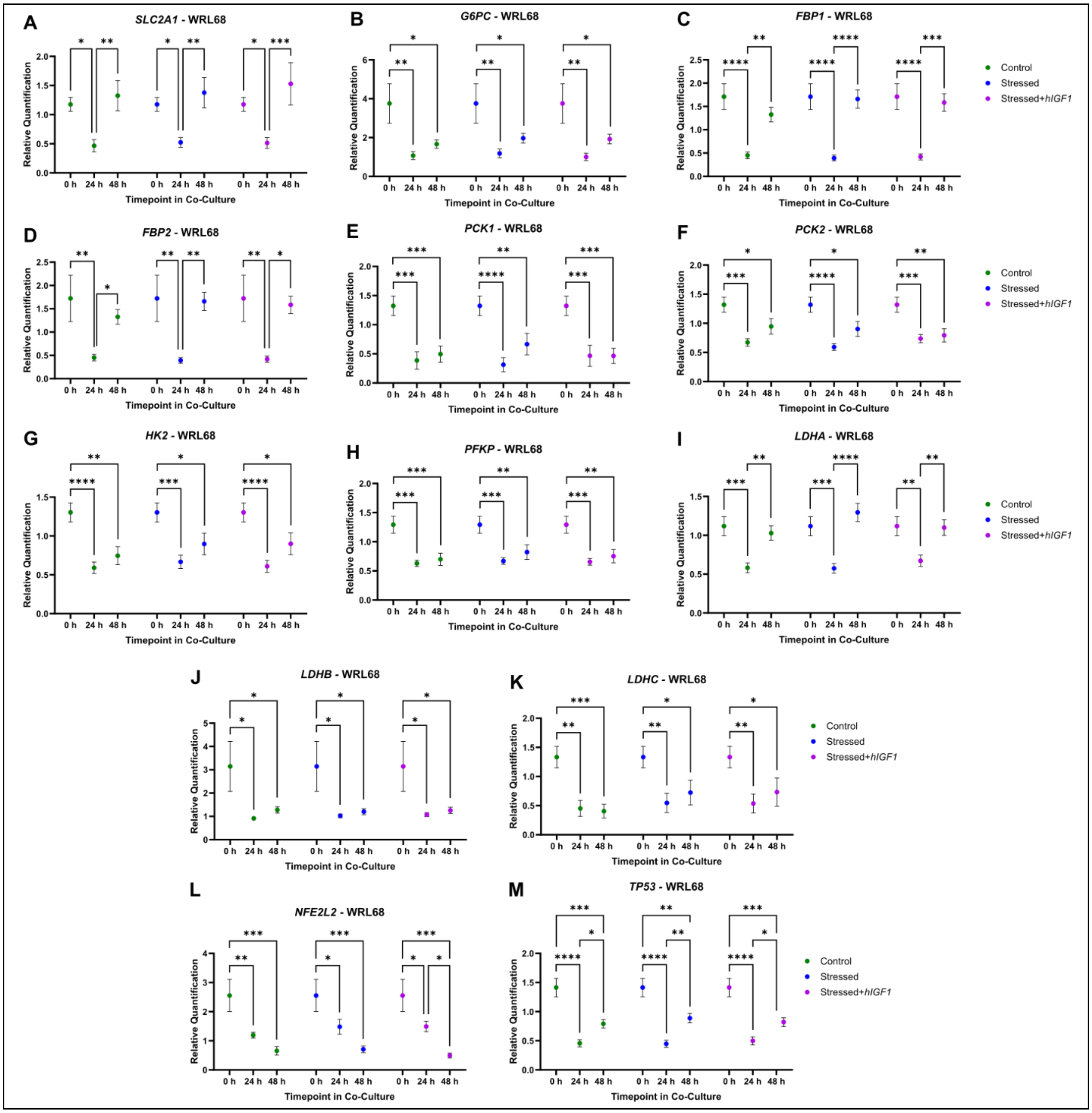


**Supplemental Figure S2. Gene expression of glucose metabolism-related factors and cell stress markers in WRL68 cells co-cultured with HPMVECs and sham-treated Control BeWo Cells (complete culture media), sham-treated Stressed BeWo cells (no FBS in culture media) or Stressed + *insulin-like 1 growth factor* (*hIGF1*) nanoparticle treated BeWo cells.** There was no difference in the gene expression of *SLC2A1* (**A**), gluconeogenesis enzymes (*G6PC* (**B**), *FBP1* (**C**), *FBP2* (**D**), *PCK1* (**E**), *PCK2* (**F**)), glycolysis enzymes (*HK2* (**G**), *PFKP* (**H**), *LDHA* (**I**), *LDHB* (**J**), *LDHC* (**K**)) nor cell stress markers (*NFE2L2* (**L**), *TP53* (**M**)) in WRL68 cells co-culture with Control, Stressed or Stressed+*hIGF1* BeWo cells**.** Data are estimated marginal mean ± SEM calculated using generalized linear modelling. n = 6 independent passages. *P<0.05; **P<0.01; ***P<0.001. HPMVEC: human placenta microvascular endothelial cells. SLC2A1: solute carrier family 2 member 1. G6PC: glucose-6-phosphatase. FBP1/2: Fructose-1,6-bisphosphatase 1/2. PCK1/2: Phosphoenolpyruvate carboxykinase 1/2. HK2: hexokinase 2. PFKP: Phosphofructokinase, platelet. LDHA/B/C: lactate dehydrogenase A/B/C. NFE2L2: Nuclear factor, erythroid 2-like 2. TP53: tumor protein 53

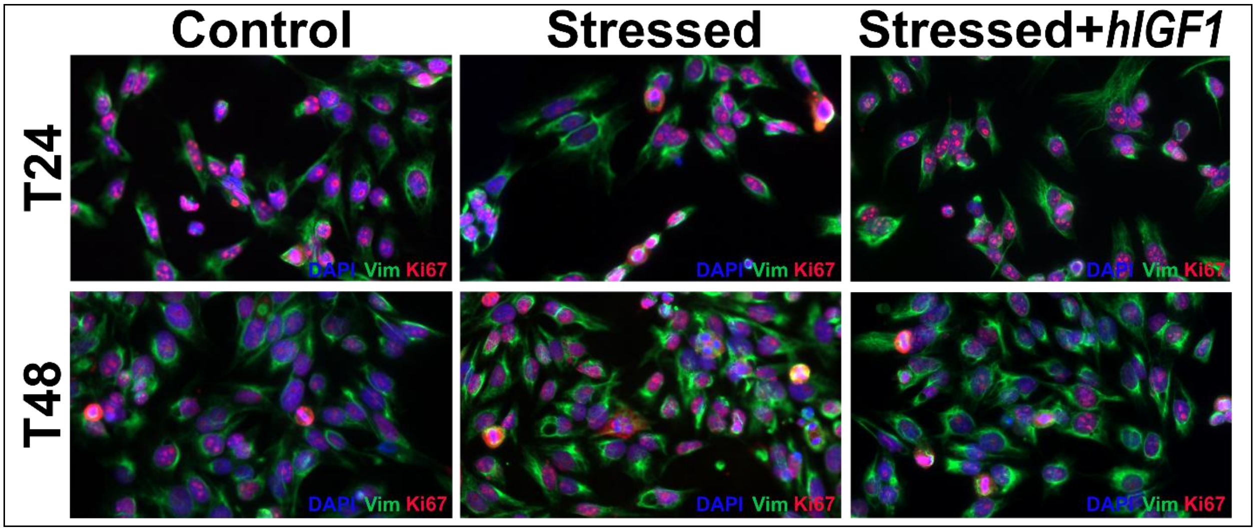


**Supplemental Figure S3. Representative images of WRL68 cells co-cultured with HPMVECs and sham-treated Control BeWo Cells (complete culture media), sham-treated Stressed BeWo cells (no FBS in culture media) or Stressed + *insulin-like 1 growth factor* (*hIGF1*) nanoparticle treated BeWo cells.** Morphologically WRL68 cells expanded and were indistinguishable when comparing co-cultured with HPMVECs and Control, Stressed or Stressed+*hIGF1* BeWo cells. n = 6 independent passages. Magnification = 40X. HPMVEC: human placenta microvascular endothelial cell. DAPI: 4',6-diamidino-2-phenylindole. Vim: Vimentin.

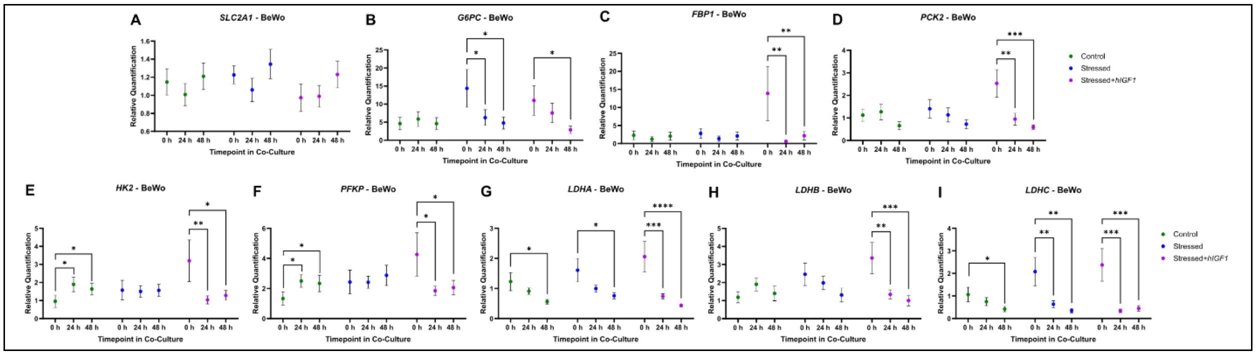


**Supplemental Figure S4. Gene expression of glucose metabolism-related factors in sham-treated Control BeWo Cells (complete culture media), sham-treated Stressed BeWo cells (no FBS in culture media) or Stressed + *insulin-like 1 growth factor* (*hIGF1*) nanoparticle treated BeWo cells co-cultured with WRL68 cells and HPMVECs.** Gene expression of *SLC2A1* (**A**), *G6PC* (**B**), *FBP1* (**C**), and *PCK2* (**D**) in Control BeWo cells did not change when co-cultured with WRL68 cells. Gene expression of *HK2* (**E**) and *PFKP* (**F**) was increased in Control BeWo cells at 24 h and remained higher at 48 h. *LDHA* (**G**) and *LDHC* (**I**) were lower in co-cultured Control BeWo cells at 48 h. In co-cultured Stressed and Stressed+*hIGF1* BeWo cells, gene expression of gluconeogenesis enzymes (**A-D**) that were higher at 0 h were reduced by 48 h, and comparable to Control BeWo cells at 48 h. Changes in gene expression of glycolysis enzymes (**E-I**) in co-cultured Stressed and Stressed+*hIGF1* BeWo cells occurred in a similar manner across the co-culture period with WRL68 cells, and were no different from Control BeWo cells at 48 h. Data are estimated marginal mean ± SEM calculated using generalized linear modelling. n = 6-12 independent passages. *P<0.05; **P<0.01; ***P<0.001. HPMVEC: human placenta microvascular endothelial cells. SLC2A1: solute carrier family 2 member 1. G6PC: glucose-6-phosphatase. FBP1: Fructose-1,6-bisphosphatase 1. PCK2: Phosphoenolpyruvate carboxykinase 2. HK2: hexokinase 2. PFKP: Phosphofructokinase, platelet. LDHA/B/C: lactate dehydrogenase A/B/C.

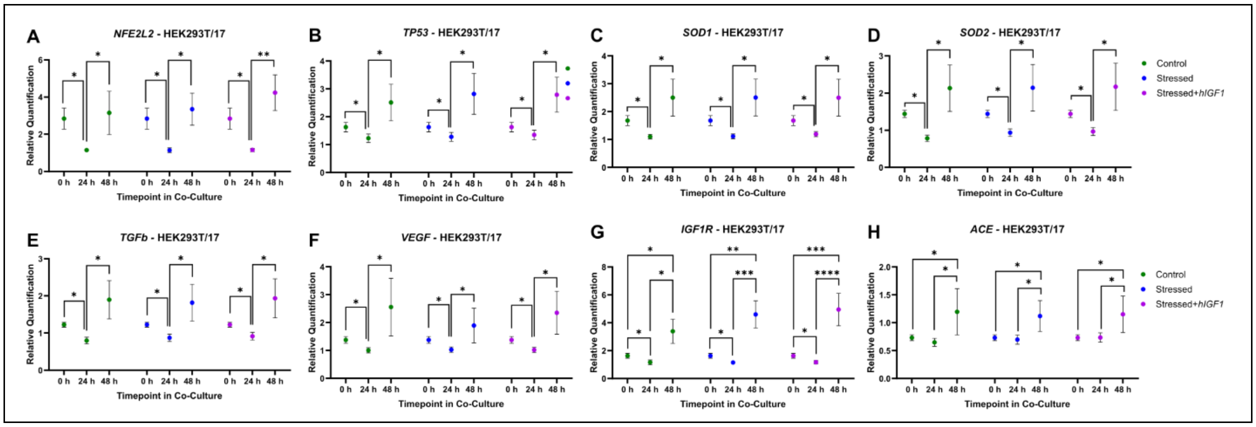


**Supplemental Figure S5. Gene expression of factors relating to epithelial cell function and blood pressure regulation in HEK293T/17 cells co-cultured with HPMVECs and sham-treated Control BeWo Cells (complete culture media), sham-treated Stressed BeWo cells (no FBS in culture media) or Stressed + *insulin-like 1 growth factor* (*hIGF1*) nanoparticle treated BeWo cells.** There was no difference in the gene expression of *NFE2L2* (**A**), *TP53* (**B**), *SOD1* (**C**), *SOD2* (**D**), *TGFb* (**E**), *VEGF* (**F**), *IGF1R* (**G**) and *ACE* (**H**) in HEK293T/17 cells co-culture with Control, Stressed or Stressed+*hIGF1* BeWo cells Data are estimated marginal mean ± SEM calculated using generalized linear modelling. n = 6 independent passages. *P<0.05; **P<0.01; ***P<0.001. HPMVEC: human placenta microvascular endothelial cells. NFE2L2: Nuclear factor, erythroid 2-like 2. TP53: tumor protein 53. SOD1/2: superoxide dismutase 1/2. TGFb: transforming growth factor beta. VEGF: vascular endothelial growth factor. IGF1R: insulin-like 1 growth factor 1 receptor. ACE: angiotensin converting enzyme

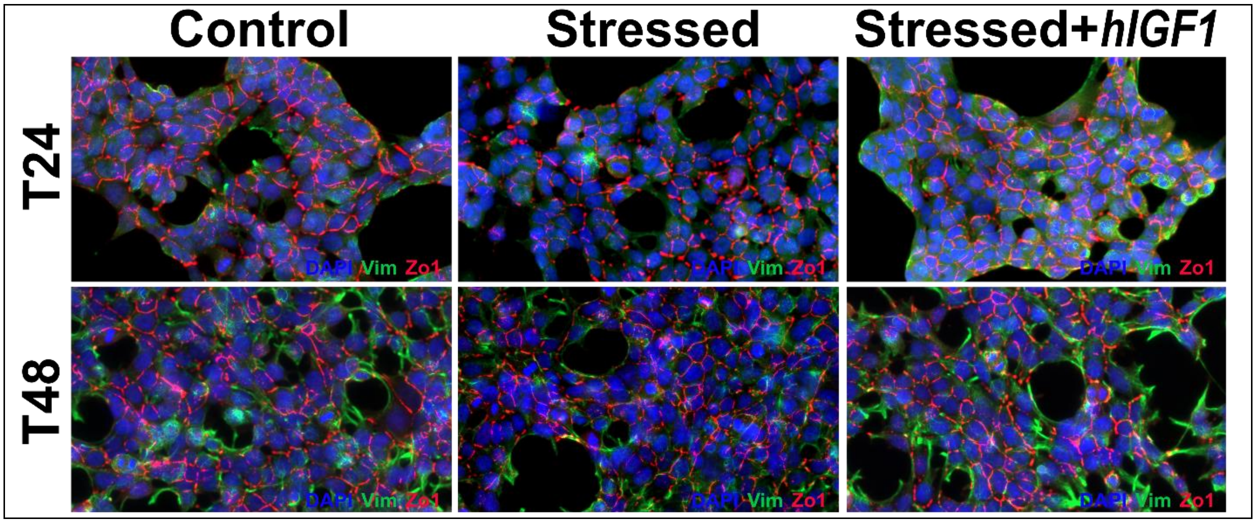


**Supplemental Figure S6. Representative images of HEK293T/17 cells co-cultured with HPMVECs and sham-treated Control BeWo Cells (complete culture media), sham-treated Stressed BeWo cells (no FBS in culture media) or Stressed + *insulin-like 1 growth factor* (*hIGF1*) nanoparticle treated BeWo cells.** Morphologically HEK293T/17 cells expanded and were indistinguishable when comparing co-cultured with HPMVECs and Control, Stressed or Stressed+*hIGF1* BeWo cells. n = 6 independent passages. Magnification = 40X. HPMVEC: human placenta microvascular endothelial cell. DAPI: 4',6-diamidino-2-phenylindole. Vim: Vimentin. Zo1: Zonula Occludens 1
